# Development of dim-light vision in the nocturnal coral reef fish family, Holocentridae

**DOI:** 10.1101/2022.05.04.490704

**Authors:** Lily G. Fogg, Fabio Cortesi, David Lecchini, Camille Gache, N. Justin Marshall, Fanny de Busserolles

## Abstract

Developmental changes to the visual systems of animals are often associated with ecological shifts. Reef fishes experience a change in habitat between larval life in the shallow open ocean to juvenile and adult life on the reef. Some species also change their lifestyle over this period and become largely nocturnal. While these ecological transitions are well documented, little is known about the ontogeny of nocturnal reef fish vision. Here, we used histology and transcriptomics to investigate visual development in 12 representative species from both subfamilies, Holocentrinae (squirrelfishes) and Myripristinae (soldierfishes), in the nocturnal coral reef fish family, Holocentridae. Results revealed that the visual systems of holocentrids are initially well-adapted to photopic conditions with pre-settlement larvae having high cone densities, high cone opsin gene expression, a broad cone opsin gene repertoire (8 genes) and a multibank retina (*i*.*e*., stacked layers of rods) comprising up to two rod banks. At reef settlement, holocentrids started to invest more in their scotopic visual system and upregulated genes involved in cell differentiation/proliferation. By adulthood, they had well-developed scotopic vision with a rod-dominated multibank retina comprising 5-17 rod banks, increased summation of rods onto ganglion cells, high rod opsin gene expression, reduced cone opsin gene expression and repertoire (1-4 genes) and upregulated phototransduction genes. Finally, although the two subfamilies shared similar ecologies across development, their visual systems diverged after settlement, with Myripristinae investing more in scotopic vision than Holocentrinae. Hence, both ecology and phylogeny likely determine the development of the holocentrid visual system.

**Summary statement:** Coral reef fishes in the family Holocentridae remodel their retina at the cellular and molecular levels to adapt to a nocturnal lifestyle during development.

## Introduction

Vision underlies many behaviours crucial to survival, most notably foraging, mating and predator avoidance (Cronin *et al*. 2014). As a result of their varied ecologies and the different light environments that they inhabit, marine fishes possess exceptionally diverse visual adaptations. These adaptations are reflected in the structure and organisation of their eye and retina (Walls 1942; Cronin 2014). The retina has four key cellular strata (in order of neural processing): the photoreceptor layer (PRL), outer nuclear layer (ONL), inner nuclear layer (INL) and ganglion cell layer (GCL) (Land and Nilsson 2012). The PRL and ONL house the outer segments (OS) and nuclei of photoreceptors, respectively, of which there are two main types: rods and cones (Lamb 2013). Rods usually contain the visual pigment, rhodopsin (RH1, rhodopsin-like middle-wavelength sensitive 1), and mediate scotopic (dim light) vision. Cones mediate photopic (bright light) and colour vision and are divided into single and double cones (*i*.*e*., two fused single cones). Single cones usually have SWS1 (short-wavelength sensitive 1) and SWS2 opsins, while double cones usually have RH2 (rhodopsin-like middle-wavelength sensitive 2) and LWS (long-wavelength sensitive) opsins [(Bowmaker 2008; Carleton *et al*. 2008; Dalton *et al*. 2017); reviewed in (Carleton *et al*. 2020)]. Each opsin class has a different range of spectral sensitivities, including UV-violet (SWS1), violet-blue (SWS2), blue-green (RH2 and RH1) and green-red (LWS), that can be tuned to match the environmental light conditions [(Lythgoe 1979; Cronin *et al*. 2014; Schweikert *et al*. 2018); reviewed in (Carleton *et al*. 2020; Musilova *et al*. 2021)].

The next retinal layer, the INL, contains the nuclei of interneurons, such as bipolar, horizontal and amacrine cells, and represents the primary stage of opponent processing for colour vision (Baden and Osorio 2019). Finally, visual signals are summated in the GCL, which sets the limits of the luminous sensitivity of the eye (*i*.*e*., more rods summating onto a single GC increases sensitivity) and spatial resolving power (*i*.*e*., acuity) (Warrant 2004). Importantly, the density and distribution of the different neural cells are usually heterogenous across the retina. Regions of the retina with high densities of a particular cell type, *i*.*e*., retinal specialisations, provide higher acuity and/or sensitivity in a specific part of an animal’s visual field (Collin 1989). Retinal specialisations often facilitate specific behavioural tasks, such as feeding or predator avoidance (Collin and Pettigrew 1988a; Luehrmann *et al*. 2020; de Busserolles *et al*. 2021).

In general, differences in the organisation and densities of the retinal cell types, the genes that they express, and their spectral sensitivities correlate well with ecological demands (Shand 1997; Stieb *et al*. 2016; Luehrmann *et al*. 2020). For instance, fishes living in dim environments (*e*.*g*., deep-sea habitat or nocturnal lifestyle) have evolved a shared array of retinal adaptations to enhance the sensitivity of their eyes, including high rod densities and low cone densities (Pankhurst 1989; Shand 1994b), high summation of rods onto ganglion cells (Shand 1994b; de Busserolles *et al*. 2020; de Busserolles *et al*. 2021), and thick PRL (with longer rods) (Wagner *et al*. 1998). Several species have pushed scotopic adaptations to an extreme level by evolving a pure rod retina (Munk 1966) or a retina with multiple layers of rods (known as a multibank retina) (McFarland 1991; de Busserolles *et al*. 2021), by duplicating their *rh1* gene (Musilova *et al*. 2019) or by combining the characteristics of both rods and cones into a single photoreceptor cell (known as transmutation) (de Busserolles *et al*. 2017). Although some of these adaptations are relatively common, little is known about their development.

During development, most marine fishes undergo significant shifts in ecology. Larval marine fishes typically inhabit the (zoo)plankton-rich upper layers of the epipelagic ocean (Job 2000; Helfman 2009), where the available light is bright and broad-spectrum ranging from UV (<400 nm) to red (>600 nm) (Boehlert 1996). However, as juveniles and adults, different species may transition to very different habitats (pelagic, estuarine, reef, deep-sea) and adopt contrasting diets (planktivory, carnivory, herbivory, corallivory), and diel activity patterns (diurnal, nocturnal, crepuscular) (King and McFarlane 2003; Helfman 2009). These changes in light environment and ecological demands are thought to be the main drivers of visual system development in marine fishes (Carleton *et al*. 2020; Musilova *et al*. 2021). Accordingly, ontogenetic changes in their visual systems have previously been correlated with changes in diet [damselfishes (Shand *et al*. 2000); surgeonfishes (Tettamanti *et al*.2019)], diel activity patterns [numerous reef fish families (Shand 1997; Siebeck and Marshall 2007)], depth [lanternfishes, hoki, hake, roughy, oreodories (Pankhurst 1987; Mas-Riera 1991)], habitat [goatfishes (Shand 1994a); mackerel icefish (Miyazaki *et al*. 2011)], body morphology [winter flounder (Evans and Fernald 1993)], and colouration [dottybacks (Cortesi *et al*. 2016)].

In fishes which adopt bright environments, visual development is characterised by typical changes in the retina. The retina is initially cone-dominated, and the densities of cones, INL cells and GC increase early in development and then decrease slightly (as retinal area expands), while rod densities undergo a minor increase (Fernald 1990; Shand 1997). These structural changes are accompanied by modifications at the molecular level, such as increased expression of cone opsin genes that are sensitive to ecologically relevant wavelengths (Shand *et al*. 2008; Cortesi *et al*. 2016; Savelli *et al*. 2018; Tettamanti *et al*. 2019). Contrastingly, in fishes which adopt dim environments, visual development seems to be characterised by a more rapid and extreme version of these changes. For example, some deep-sea fishes seem to possess cones as larvae but progress to having only rods in adulthood (Bozzano *et al*. 2007; de Busserolles *et al*. 2014a; Lupše *et al*. 2021). However, most of the previous studies on visual development in dim environments focused on deep-sea fishes (Cortesi *et al*. 2020). Conversely, very little is known about how the visual system develops in nocturnal reef fishes [but see (Shand 1997; Job and Shand 2001)].

Here, we investigated visual development in the nocturnal reef fish family, Holocentridae. Larvae from both subfamilies, Holocentrinae (squirrelfishes) and Myripristinae (soldierfishes), inhabit the upper pelagic ocean where they feed on zooplankton (Tyler *et al*. 1993; Sampey *et al*. 2007). Following metamorphosis, most holocentrids (except for a few species that migrate to deeper waters) migrate to a shallow tropical coral reef habitat (Nelson 1994) and adopt a nocturnal lifestyle feeding on benthic crustaceans (Holocentrinae) or pelagic zooplankton (Myripristinae) (Gladfelter and Johnson 1983; Greenfield 2002; Greenfield *et al*. 2017). We recently characterised the visual systems of adult holocentrids showing that they possess well-developed scotopic vision with a rod-dominated retina arranged into multiple banks (de Busserolles *et al*. 2021). The complexity of their multibank retina resembles that of some deep-sea fishes, with up to 7 and 17 banks in Holocentrinae and Myripristinae, respectively (de Busserolles *et al*. 2021). Furthermore, their rod spectral sensitivities have been shown to be tuned to their preferred depth, *i*.*e*., blue-shifted when living deeper compared to red-shifted in the shallows (Toller 1996; Yokoyama and Takenaka 2004). Adults also have some degree of photopic vision, more so in Holocentrinae than Myripristinae, with the presence of single cones expressing *sws2*, and double cones expressing 1-2 *rh2*s, all well organised into retinal specialisations (de Busserolles *et al*. 2021).

Although the visual systems of adult holocentrids have been well-characterised, little is known about how they develop. Thus, we used a multidisciplinary approach to examine the anatomical structure, cell densities, opsin gene expression and whole-transcriptome differential gene expression in the retina at key ontogenetic stages (pre-settlement larvae, settlement larvae, settled juveniles and adults). We studied shallow reef-dwelling species from three genera (*Sargocentron, Neoniphon* and *Myripristis*) spanning both holocentrid subfamilies, as well as an adult for a deeper-dwelling species (*Ostichthys* sp.). Using this approach, we addressed the following aims: 1) to assess how the holocentrid visual system changes as they shift from a bright to a dim environment, and 2) to study the development of their deep-sea-like multibank retina.

## Materials and methods

### Animal collection and retinal tissue preservation

Details of all animals used in this study are given in Table S1. Pre-settlement larvae, which are pelagic larvae close to transitioning to the reef, were collected on the Great Barrier Reef around Lizard Island, Australia (14°40′S, 145°27′E) using light traps in February 2020. Settlement larvae, larvae that have just transitioned to the reef, were collected using a crest net positioned on the reef crest of the lagoon at Temae, Moorea, French Polynesia (17°29’S, 149°45’W) in February and March 2019 (Lecchini *et al*. 2004; Besson *et al*. 2017). Settled juveniles were larvae caught in light traps on Lizard Island which were allowed to metamorphose and further develop for two weeks in outdoor aquaria exposed to natural light in March 2017. Adults were collected with either spearguns or pole and lines on the reefs surrounding Moorea in March 2018 and 2019 or collected with clove oil and hand nets at Lizard Island in February 2020. Some adults were also sourced from a supplier, Cairns Marine (Cairns Marine Pty Ltd, Cairns, Australia; https://www.cairnsmarine.com/), that collects from the northern Great Barrier Reef.

Fish collection and euthanasia followed procedures approved by the University of Queensland Animal Ethics Committee (QBI 304/16). All collections within Australia were conducted under a Great Barrier Reef Marine Park Permit (G17/38160.1) and Queensland General Fisheries Permit (180731) and all collections in French Polynesia were conducted in accordance with French regulations. Following euthanasia, all individuals were photographed with a ruler and their body size [total length (TL) and standard length (SL)] and eye diameter subsequently measured from photographs using Fiji v1.53c [National Institutes of Health, USA; (Schindelin *et al*. 2012)]. Eyes were immediately enucleated, the cornea and lens removed, and the eye cup preserved in either RNAlater (Sigma-Aldrich, R0901) or 4% paraformaldehyde [PFA; 4% (w/v) PFA in 0.01M phosphate-buffered saline (PBS), pH 7.4] depending on the analysis (see below for details).

### Histology

Retinal histology was conducted on PFA-fixed eyes from the following individuals: three pre-settlement larvae (*Sargocentron rubrum, n*=3), four settlement larvae (*Myripristis berndti, n*=1; *M. kuntee, n*=3), two settled juveniles (*S. rubrum, n*=2) and nine adults (*S. rubrum, n*=3; *S. microstoma, n=1; M. berndti, n*=2; *M. kuntee, n*=1; *M. violacea, n*=1; *Ostichthys* sp., *n*=1). To account for intraretinal variability (de Busserolles *et al*. 2021), two (dorsal and ventral) or five (dorsal, ventral, central, nasal and temporal) retinal regions were sampled for pre-settlement larvae and later stages, respectively. Notably, for the *Ostichthys* sp., tissue quality was only sufficient to examine one region (ventral). Briefly, a small square of retina was dissected from each region, post-fixed in 2.5% glutaraldehyde and 2% osmium tetroxide, dehydrated in ethanol and/or acetone, and embedded in EPON resin (ProSciTech). All tissue processing was done in a BioWave Pro tissue processor (PELCO).

Radial 1 μm-thick retinal sections were cut with a glass knife on a Leica ultramicrotome (Ultracut UC6) and stained with a solution of 0.5% toluidine blue and 0.5% borax. Retinal sections were viewed with a 63X objective (oil, 1.4 numerical aperture, 0.19 mm working distance, 0.102 μm/pixel) on a Zeiss Axio upright microscope (Imager Z1) and brightfield images acquired with Zeiss Axiocam 506 mono and 512 colour microscope cameras. Rod outer segment (ROS) length, PRL thickness, and whole retinal thickness were then measured from micrographs using Fiji. A body size range at which the full complement of banks (*i*.*e*., the maximum number of rod banks detected across all individuals/stages for each species) was reached was determined by comparing TL in individuals with the full complement to the maximal TL published on FishBase (https://www.fishbase.se) (Froese and Pauly 2019).

### Cell density estimations

Retinal cell densities were estimated from transverse retinal sections, a method widely employed for marine teleosts for over 50 years (Munk 1965; Locket 1980; Shand 1997; Taylor *et al*. 2015). Cell densities were compared at different stages in the same species for Holocentrinae (*S. rubrum*), and in two species in the same genus for Myripristinae due to a limitation in the number of specimens at specific stages (settlement: *M. kuntee*, adult: *M. berndt*i). However, to make sure that the data were comparable between the different species from Myripristinae, the densities in settlement *M. kuntee* were also compared to one settlement *M. berndti* and the densities in adult *M. berndti* were compared to one adult *M. kuntee* (Fig. S1). Briefly, using Fiji, images were cropped to obtain retinal strips of 250 μm (horizontal length) for lower-density cell types (*i*.*e*., cone and ganglion cells) in adults, 100 μm for lower-density cell types in larvae, and 40 μm for higher-density cell types (*i*.*e*., ONL and INL cells) for all life stages. These counting frames were optimised by conducting trials with several frame sizes and taking the minimum frame size that produced counts ≥95% similar to those attained with the largest frame (assumed to be the most accurate). The number of cone OS, ONL nuclei, INL nuclei and GCL nuclei were counted for three sections per sample using the cell counter plugin in Fiji. Subsequently, counts were corrected for section thickness using Abercrombie’s correction (Abercrombie 1946) and the density of each retinal cell type per 0.01 mm^2^ of retina was calculated. Rod densities were calculated as the difference between ONL nuclei and cone OS densities (Shand 1994a). Graphs throughout the manuscript were generated using GraphPad Prism software v8.3.1 (www.graphpad.com).

### Transcriptome sequencing, quality control and de novo assembly

Retinal transcriptomes were sequenced for a total of 22 individuals in Holocentridae: two pre-settlement larvae (*S. rubrum, n=*2), seven settlement larvae (*S. punctatissimum, n*=4; *M. berndti, n*=1; *M. pralinia, n*=1; *M. kuntee, n*=1), six settled juveniles (*S. rubrum, n*=3; *S. cornutum, n*=1; *Neoniphon sammara, n*=2) and seven adults (*S. punctatissimum, n*=3; *S. rubrum, n*=2; *M. kuntee, n*=1; *Ostichthys* sp., *n*=1). The adult dataset was completed with previously published transcriptomic (*S. rubrum, n*=1; *S. spiniferum, n*=1; *S. diadema, n*=1; *N. sammara, n*=3; *M. berndti, n*=4; *M. jacobus, n*=2, *M. murdjan, n*=1) and genomic (*M. jacobus, n*=1) data (Malmstrøm *et al*. 2017; Musilova *et al*. 2019; de Busserolles *et al*. 2021), resulting in a total dataset of 35 retinal transcriptomes and one genome spanning 12 species.

Briefly, initial retinal tissue digestion was conducted using Proteinase K (New England Biolabs). Total RNA was extracted and isolated from the retinas using the Monarch Total RNA Miniprep Kit (New England Biolabs) and genomic DNA was removed using RNase-free DNase (Monarch). Quality and yield of isolated RNA was assessed using the Eukaryotic Total RNA 6000 Nano kit (Agilent technologies) and the Queensland Brain Institute’s Bioanalyser 2100 (Agilent technologies). RNA extractions were shipped on dry ice and whole-retina transcriptome libraries were prepared from total RNA using the NEBNext Ultra RNA library preparation kit for Illumina (New England Biolabs) at Novogene’s sequencing facilities in Beijing, Hong Kong, and Singapore. The concentration of each library was checked using a Qubit dsDNA BR Assay kit (ThermoFisher) prior to barcoding and pooling at equimolar ratios. Libraries were sequenced as 150 bp paired-end reads on a HiSeq 2500 using Illumina’s SBS chemistry version 4. Libraries were trimmed and *de novo* assembled as described in de Busserolles *et al*. (2017). Briefly, read quality was assessed using FastQC (v0.72), raw reads were trimmed and filtered using Trimmomatic (v0.36.6) and transcriptomes were *de novo* assembled with Trinity (v2.8.4) using the genomics virtual laboratory on the Galaxy platform [usegalaxy.org; (Afgan *et al*. 2018)].

### Opsin gene mining, phylogenetic reconstruction and expression analyses

All cytochrome C oxidase subunit I (COI) and opsin gene extractions and expression analyses were conducted in Geneious Prime v2021.1.1 (Biomatters Ltd). Initially, COI genes were extracted from *de novo* assembled transcriptomes for species identification by mapping to species-specific references from Genbank (https://www.ncbi.nlm.nih.gov/genbank/) with medium sensitivity settings. Opsin gene extractions were performed by mapping assembled transcriptomes to published opsin coding sequences (CDS) of the dusky dottyback (*Pseudochromis fuscus)* (Cortesi *et al*. 2016) with customised sensitivity settings (fine tuning, none; maximum gap per read, 15%; word length, 14; maximum mismatches per read, 40%; maximum gap size, 50 bp; index word length, 12; paired reads must both map). Contigs mapped to COI and opsin references were scored for similarity against publicly available sequences using BLASTn (NCBI, Bethesda, MD, https://blast.ncbi.nlm.nih.gov/Blast.cgi). One of the limitations of *de novo* assembly of highly similar genes (such as opsin gene paralogs) using short-read transcripts is that it can produce erroneous (chimeric) sequences or fail to reconstruct lowly expressed transcripts. Thus, a second approach was also employed using a manual extraction method from back-mapped reads to verify the initially extracted opsin genes, as per de Busserolles *et al*. (2017).

During manual gene extraction, filtered paired reads were mapped against *P. fuscus* reference opsin CDS (with previously stated customised sensitivity settings) in Geneious Prime. Matching reads were connected by following single nucleotide polymorphisms (SNPs) across genes with continual visual inspection for ambiguity and were extracted as paired mates to mitigate sequence gaps. The consensus of an assembly of these extracted reads was used as the reference for low sensitivity (high accuracy, 100% identity threshold) mapping. Partial CDS extractions were cyclically mapped using the low sensitivity approach to prolong and subsequently remap reads until a complete CDS was obtained.

To confirm the identity of each gene, full coding sequences were preliminarily checked on BLASTn and then used in conjunction with a reference dataset obtained from Genbank (www.ncbi.nlm.nih.gov/genbank/) and Ensembl (www.ensembl.org/) to phylogenetically reconstruct the opsin gene phylogeny (de Busserolles *et al*. 2017; de Busserolles *et al*. 2021). All opsin gene sequences were aligned using the MUSCLE plugin v3.8.425 (Edgar 2004) in Geneious Prime. MrBayes v3.2.6 (Ronquist *et al*. 2012) on CIPRES (Miller *et al*. 2010) was used to reconstruct a phylogenetic tree from the aligned sequences using the following parameters: GTR+I+G model, two independent MCMC searches with four chains each, 10 million generations per run, 1000 generations sample frequency, and 25% burn-in. The generated tree was edited in Figtree v1.4.4 (Rambaut and Drummond 2012).

For differential expression analyses, opsin gene paralogs were first scored on similarity using pairwise/multiple alignments. The similarity score minus one was used as the gene-specific maximum % mismatch threshold for mapping (paired) transcripts back onto complete extracted opsin CDS to ensure that reads did not map to multiple paralogs. The proportional expression of rod vs. cone opsin genes was calculated as the number of reads mapped to either *rh1* or all cone genes divided by the number of reads mapped to all genes, adjusted to account for differing gene lengths. The proportional expression of single vs. double cone opsin genes was calculated as the number of reads mapped to each single (*i*.*e*., *sws1* and *sws2*) or double (*i*.*e*., *rh2* and *lws*) cone opsin gene copy divided by the number of reads mapped to all single or double cone opsin genes, respectively, and adjusted for gene length.

### Whole-transcriptome differential gene expression analyses

Further analyses were conducted to search for differentially expressed genes (DEGs) over development across the whole retinal transcriptome for *S. punctatissimum* (settlement larvae, *n*=3; adults, *n*=3). To control for diel fluctuations in visual gene expression, all individuals were euthanised at 8:30 am (Yourick *et al*. 2019). All analyses were conducted using the genomics virtual laboratory on the Galaxy platform. Firstly, the quality of the sequencing reads was assessed using FastQC (v0.72) and raw reads were subsequently trimmed and filtered using Trimmomatic (v0.36.6) as described previously. Given that a high-quality reference genome was not available for the species sequenced, a reference transcriptome was *de novo* assembled using Trinity (v2.8.4) from the combined reads of one paired-end library from each life stage (with settings described above). A mapping-based estimation of transcript abundance was then obtained using Salmon quant v0.14.1.2 (using default settings except specifying paired-end, stranded reads (SF), discarding orphan quasi, validating mappings and mimicking Bowtie) (Patro *et al*. 2017). Differentially expressed transcripts were identified using DEseq2 v2.11.40.6 (using default settings with the following changes: setting input data to Transcripts Per Kilobase Million (TPM) values generated by Salmon, outputting normalised counts table and no independent filtering) (Love *et al*. 2014). The DEseq2 result file was filtered (Filter v 1.1.0) on the adjusted p-value column (≤ 0.05) to obtain DEGs.

Separate lists of up- and down-regulated DEGs were obtained by filtering for positive and negative values in the log fold change column of the DEseq2 result file. These lists were re-formatted (using FASTA-to-TABULAR v1.1.1, Cut v1.0.2, Sort v1.1.1, Join v1.1.2 and TABULAR-to-FASTA v1.1.1 tools) to generate simple gene ID-to-sequence lists of DEGs in FASTA format. The top BLAST hit from the UniProtKB/Swiss-Prot database (The UniProt Consortium 2018) was obtained using NCBI BLAST+ blastx v0.3.3 (using default settings but selecting blastx analysis) and Blast top hit descriptions v0.0.9, and was used to annotate DEG sequences (Altschul *et al*. 1997; Camacho *et al*. 2009; Cock *et al*. 2013; Cock *et al*. 2015). For the top ten DEGs, QuickGO was used to manually search for gene ontology (GO)/function (Binns *et al*. 2009). For the whole dataset, PANTHER was used via The Gene Ontology Resource website to perform a GO enrichment analysis for biological processes (Ashburner *et al*. 2000; Gene Ontology Consortium 2021). PANTHER analyses used *Oryzias latipes* as the reference, used Fisher’s exact test, calculated false discovery rate (FDR), used an FDR-adjusted p-value of < 0.01 and filtered results for GO terms with fold enrichment ≥2.

## Results

### Multibank retina structure

Retinal sections were taken at different life stages from species in each subfamily in Holocentridae to assess the structure of their multibank retina. In all species and stages, rods were arranged in banks in at least part of the retina. Moreover, rod banking increased with size/age. Pre-settlement larvae of *S. rubrum* (the only species obtained at this stage) had two rod banks in the dorsal retina but only one bank in the ventral retina (Fig. 1C). In settled juveniles, this increased to 3-4 rod banks in all regions and in adults, increased to five banks in the dorsal, nasal and ventral regions and 7 banks in the temporal and central retina (Fig. 1A-E, Fig. S2). Settlement larvae of *M. kuntee* and *M. berndti* had 3-4 rod banks in all regions, while adults possessed 12-13 banks in all regions except the ventral retina, which had 17 banks (Fig. 1F-I, Fig. S2). Finally, the adult specimen of the deeper dwelling soldierfish, *Ostichthys* sp., had approximately 10 rod banks in the ventral retina (Fig. S3). Across the family, the full complement of banks was attained by the time fish reached 40-60% of maximal size.

**Fig. 1.**
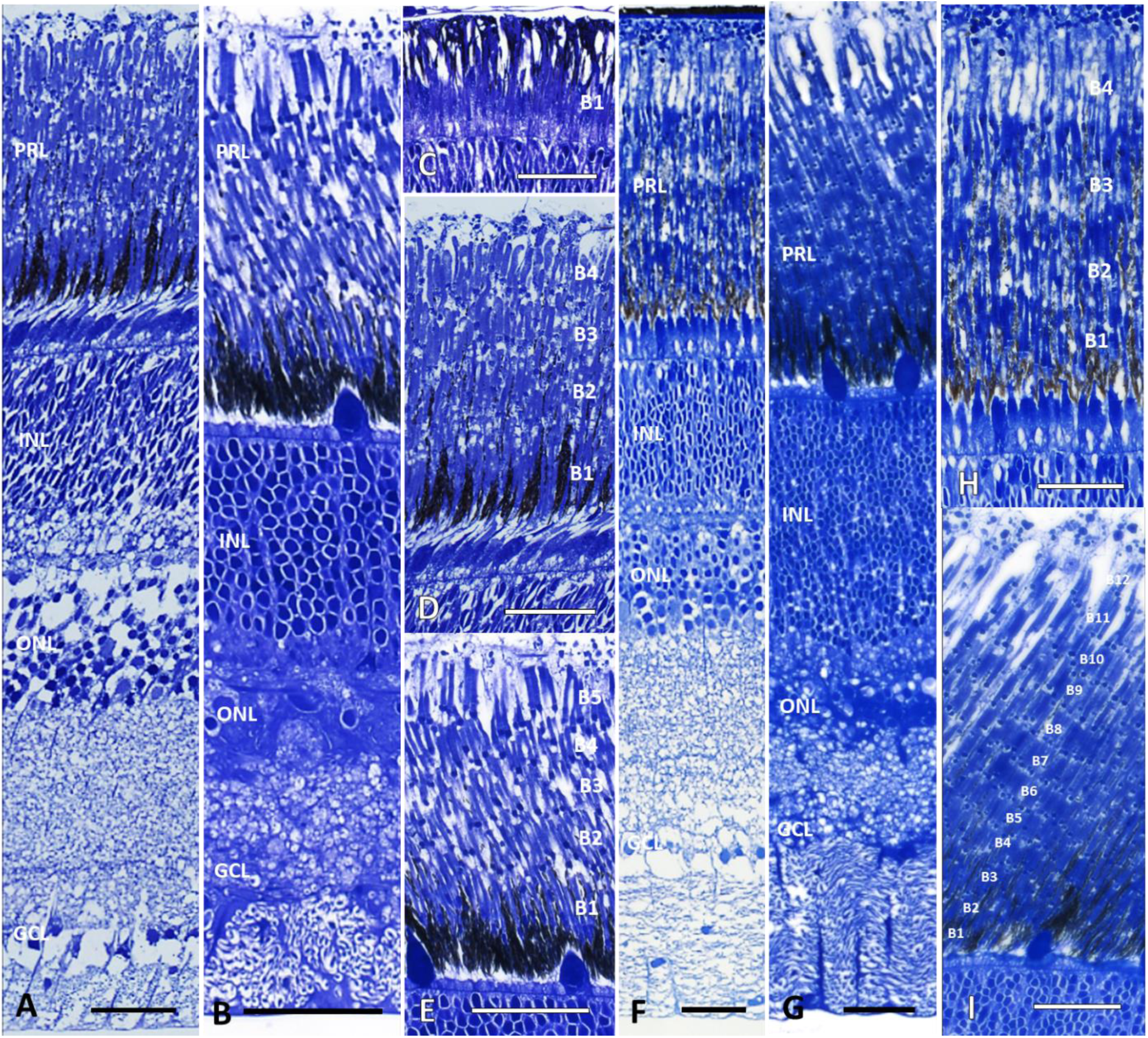
The multibank retina of holocentrids over ontogeny. Representative radial sections of the entire retina (A, B, F, G) or the photoreceptor layer (C-E, H, I) for species in Holocentridae. Rod banks in the photoreceptor layer are numbered as B_1-n_. (A-E) Holocentrinae: pre-settlement larval *Sargocentron rubrum* (C), settled juvenile *S. rubrum* (A, D) and adult *S. microstoma* (B, E). (F-I) Myripristinae: settlement-stage *Myripristis kuntee* (F, H) and adult *M. violacea* (G, I). PRL, photoreceptor layer; INL, inner nuclear layer; ONL, outer nuclear layer; GCL, ganglion cell layer. Scale bars: 30 µm. Sections for all retinal regions are given in Fig. S2.

In species from both subfamilies (Holocentrinae: *S. microstoma;* Myripristinae: *M. berndti*), the addition of rod banks between settlement larvae/settled juveniles and adults resulted in an increase in the PRL thickness (Table S2). The regions with the greatest increase in rod banks over ontogeny showed the greatest increase in PRL thickness and peak PRL thickness matched peak rod banking in adults. However, the ontogenetic increase in rod banking did not result in a linear increase in PRL thickness due to concurrent shortening of the ROS with age (Table S2).

### Retinal cell densities

The densities of different retinal cell types (rods, cones, INL cells and GC) were estimated in different regions of the retina for species in Holocentrinae (*S. rubrum*) and Myripristinae (*M. kuntee, M. berndti*) (Fig. 2). In *S. rubrum*, between pre-settlement larval and settled juvenile stages, mean cone, INL and GC densities decreased across the retina, by 65-75%, 19-42% and 31-39%, respectively (% range for the different regions) (Fig. 2, Table 2). Concurrently, rod densities and rod:GC summation increased in all regions, by 34-63% and 120-136%, respectively. Between settled juvenile and adult stages, cone, INL and GC densities continued to decrease across the retina, by 81-92%, 77-90% and 83-95%, respectively. Concurrently, rod densities and rod:GC summation further increased by 10-44% and 663-2,073%, respectively.

**Table 1.** Retinal cell measurements used for Abercrombie’s correction. Correction factors are mean (in µm) of six measurements per cell type and individual for the dorsal, ventral, central, nasal and temporal retina in Holocentrinae and Myripristinae (see legend of Fig. 2 for number of individuals). GC, ganglion cell; INL, inner nuclear layer; ONL, outer nuclear layer; OS, outer segment.

|  |  | *Sargocentron rubrum* |  |  |  |
| --- | --- | --- | --- | --- | --- |
|  |  | ONL nuclei | Cone OS | INL nuclei | GC nuclei |
| Pre-settlement larvae | Dorsal | 2.6 | 4.2 | 4.2 | 6.2 |
|  | Ventral | 2.5 | 4.2 | 4.3 | 5.9 |
| Settled juvenile | Dorsal | 2.8 | 7.9 | 4.5 | 5.3 |
|  | Ventral | 2.6 | 7.4 | 4.2 | 5.7 |
|  | Central | 2.7 | 7.8 | 4.4 | 5.3 |
|  | Nasal | 2.7 | 8 | 4.7 | 6.1 |
|  | Temporal | 2.5 | 6.9 | 4.3 | 5.4 |
| Adult | Dorsal | 2.7 | 12.5 | 5 | 6.6 |
|  | Ventral | 2.8 | 11.7 | 5.1 | 5.6 |
|  | Central | 2.8 | 11.4 | 5.1 | 5.9 |
|  | Nasal | 2.6 | 11.6 | 4.7 | 5.6 |
|  | Temporal | 2.5 | 10.9 | 5.1 | 5.8 |
|  |  | *Myripristis* spp. |  |  |  |
|  |  | ONL nuclei | Cone OS | INL nuclei | GC nuclei |
| Settlement larvae | Dorsal | 2.8 | 5.5 | 4.8 | 6 |
|  | Ventral | 2.9 | 5.5 | 4.9 | 7 |
|  | Central | 2.9 | 5.2 | 5.5 | 7 |
|  | Nasal | 2.8 | 4.9 | 5.5 | 6.9 |
|  | Temporal | 2.7 | 5 | 4.9 | 6.5 |
| Adult | Dorsal | 2.5 | 12.1 | 5 | 7.1 |
|  | Ventral | 2.7 | 11.1 | 5.4 | 6.6 |
|  | Central | 2.9 | 10.8 | 5.1 | 6.9 |
|  | Nasal | 2.7 | 12.3 | 5 | 7.4 |
|  | Temporal | 2.6 | 12.9 | 5.1 | 5.8 |

**Table 2.** Retinal cell densities in holocentrids over ontogeny. Abercrombie-corrected retinal cell densities in different life stages/species in Holocentrinae and Myripristinae (see legend of Fig. 2 for number of individuals). Values are given mean ± s.e.m. in µm for each retinal region (*i*.*e*., dorsal, ventral, central, nasal or temporal). Dorsal and ventral regions were sampled for pre-settlement larvae (due to eye size) and other regions are marked as n.a. Abbreviations: INL, inner nuclear layer; GC, ganglion cells.

|  |  | Pre-settlement larvae - *S. rubrum* |  | Settled juveniles – *S. rubrum* |  | Adults – *S. rubrum* |  |
| --- | --- | --- | --- | --- | --- | --- | --- |
| Cell type | Region | Mean | s.e.m. | Mean | s.e.m. | Mean | s.e.m. |
| Rods | Dorsal | 4447.7 | 570.8 | 7261.9 | 706.9 | 7982.2 | 1714.3 |
|  | Ventral | 3853.8 | 342.6 | 5149.1 | 139.6 | 6248.3 | 612.5 |
|  | Central | n.a. | n.a. | 7225.5 | 1444.3 | 8963.3 | 753.4 |
|  | Nasal | n.a. | n.a. | 6796.4 | 1095.6 | 8038.0 | 1664.9 |
|  | Temporal | n.a. | n.a. | 6672.0 | 283.8 | 9599.0 | 477.0 |
| Cones | Dorsal | 528.9 | 102.3 | 127.4 | 6.3 | 10.3 | 1.7 |
|  | Ventral | 459.4 | 19.1 | 158.2 | 26.2 | 18.4 | 1.0 |
|  | Central | n.a. | n.a. | 137.5 | 11.6 | 23.5 | 3.6 |
|  | Nasal | n.a. | n.a. | 149.9 | 4.5 | 22.3 | 4.3 |
|  | Temporal | n.a. | n.a. | 198.3 | 42.4 | 38.0 | 3.2 |
| INL cells | Dorsal | 2678.1 | 361.9 | 1558.7 | 203.3 | 285.1 | 43.6 |
|  | Ventral | 3091.3 | 501.4 | 2514.5 | 31.5 | 247.5 | 48.6 |
|  | Central | n.a. | n.a. | 1971.4 | 210.4 | 447.9 | 91.6 |
|  | Nasal | n.a. | n.a. | 2084.4 | 242.3 | 388.5 | 98.1 |
|  | Temporal | n.a. | n.a. | 2802.8 | 206.6 | 605.3 | 27.9 |
| GC | Dorsal | 251.5 | 19.8 | 172.8 | 6.1 | 9.9 | 2.5 |
|  | Ventral | 267.2 | 2.2 | 162.4 | 11.9 | 8.3 | 1.4 |
|  | Central | n.a. | n.a. | 183.0 | 17.1 | 27.6 | 4.5 |
|  | Nasal | n.a. | n.a. | 158.4 | 0.5 | 16.8 | 5.1 |
|  | Temporal | n.a. | n.a. | 278.4 | 6.5 | 48.7 | 6.0 |
| Rod:GC | Dorsal | 17.7 | 1.9 | 41.9 | 5.3 | 909.3 | 271.8 |
|  | Ventral | 14.4 | 1.4 | 31.8 | 1.5 | 775.1 | 77.2 |
|  | Central | n.a. | n.a. | 38.8 | 3.9 | 345.9 | 74.5 |
|  | Nasal | n.a. | n.a. | 43.0 | 7.1 | 526.1 | 24.2 |
|  | Temporal | n.a. | n.a. | 24.0 | 0.4 | 182.6 | 16.1 |
|  |  | Settlement larvae – *M. kuntee* |  | Adults – *M. berndti* |  |  |  |
| Cell type | Region | Mean | s.e.m. | Mean | s.e.m. |  |  |
| Rods | Dorsal | 6992.8 | 391.8 | 17266.5 | 3427.1 |  |  |
|  | Ventral | 6590.9 | 373.6 | 25986.9 | 587.2 |  |  |
|  | Central | 7070.6 | 1123.4 | 14430.2 | 1207.2 |  |  |
|  | Nasal | 6330.3 | 722.9 | 15937.6 | 190.8 |  |  |
|  | Temporal | 4898.1 | 596.1 | 15452.5 | 2060.8 |  |  |
| Cones | Dorsal | 200.7 | 6.0 | 7.2 | 0.4 |  |  |
|  | Ventral | 151.8 | 14.8 | 7.1 | 1.4 |  |  |
|  | Central | 157.8 | 7.7 | 12.4 | 2.6 |  |  |
|  | Nasal | 198.0 | 15.8 | 9.6 | 0.6 |  |  |
|  | Temporal | 220.3 | 20.8 | 9.2 | 1.7 |  |  |
| INL cells | Dorsal | 1971.5 | 42.4 | 437.9 | 8.5 |  |  |
|  | Ventral | 2408.7 | 306.5 | 494.8 | 0.0 |  |  |
|  | Central | 1696.6 | 52.4 | 334.6 | 1.3 |  |  |
|  | Nasal | 1884.1 | 228.2 | 430.7 | 49.1 |  |  |
|  | Temporal | 3270.1 | 220.7 | 340.4 | 4.0 |  |  |
| GC | Dorsal | 107.3 | 2.1 | 5.9 | 1.0 |  |  |
|  | Ventral | 116.2 | 6.3 | 7.0 | 0.0 |  |  |
|  | Central | 138.6 | 4.5 | 12.6 | 0.2 |  |  |
|  | Nasal | 109.3 | 5.7 | 7.3 | 1.2 |  |  |
|  | Temporal | 184.9 | 44.0 | 10.0 | 2.5 |  |  |
| Rod:GC | Dorsal | 65.3 | 4.8 | 2934.4 | 105.1 |  |  |
|  | Ventral | 56.8 | 2.9 | 3803.4 | 0.0 |  |  |
|  | Central | 51.4 | 9.3 | 1141.8 | 75.7 |  |  |
|  | Nasal | 58.6 | 8.5 | 2253.0 | 408.2 |  |  |
|  | Temporal | 30.6 | 8.7 | 1700.1 | 620.4 |  |  |

**Fig. 2.**
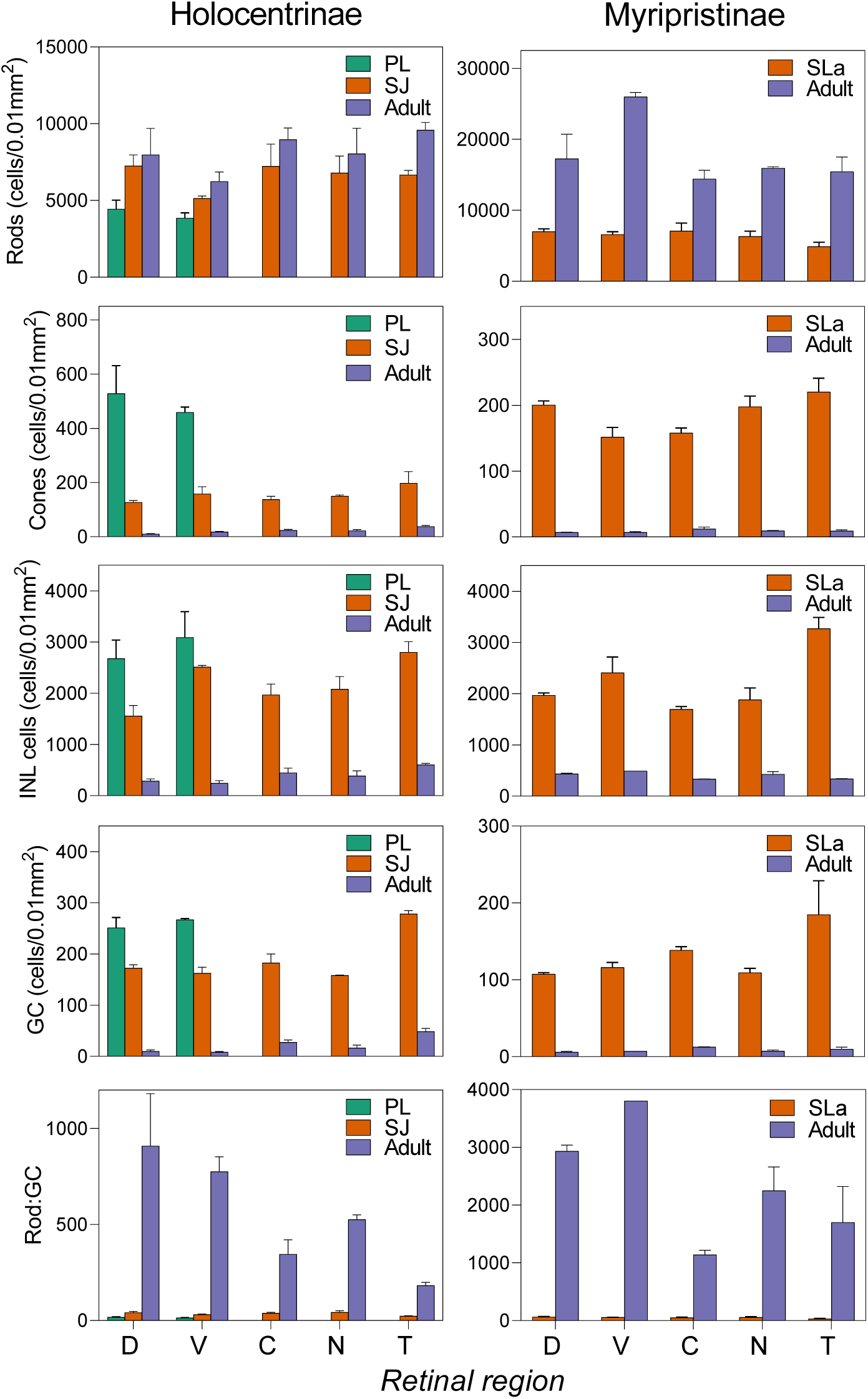
Retinal cell densities in holocentrids over ontogeny. Abercrombie-corrected densities of rods, cones, inner nuclear layer (INL) cells and ganglion cells (GC), and rod:GC summation in the dorsal (D), ventral (V), central (C), nasal (N) and temporal (T) retina in Holocentrinae [*Sargocentron rubrum* pre-settlement larvae (*n*=3), settled juveniles (*n*=2) and adults (*n*=3)] and Myripristinae [*Myripristis kuntee* settlement larvae (*n*=3) and *M. berndti* adults (*n*=2)]. Cell densities are cells per 0.01 mm^2^ of retina given as mean ± s.e.m. Green, pre-settlement larvae (PL); orange, settlement larvae (SLa; Myripristinae) or settled juveniles (SJ; Holocentrinae); purple, adults. Cell measurements used for Abercrombie’s correction are given in Table 1.

A similar developmental pattern was observed in *Myripristis* spp. (Fig. 2, Table 2). Since cell densities were found to be similar between *M. kuntee* and *M. berndti* (Fig. S1), a comparison between stages was done using the two species to increase sample size. Between settlement and adulthood, cone, INL cell and GC densities decreased across the retina by 92-96%, 77-90% and 90-95%, respectively. Concurrently, rod densities and rod:GC summation increased by 104-294% and 2,123-6,592%, respectively.

Intraretinal shifts in peak cell densities were also found in all holocentrids examined (Fig. 2, Table 2). Around settlement, all species had higher cone, INL cell and GC densities in the temporal retina. At adulthood, *S. rubrum* retained these peak densities in the temporal retina, while *Myripristis* spp. shifted its peak cone and GC densities centrally, and its peak INL cell densities ventrally. Conversely, rod densities did not peak in the same regions for *S. rubrum* and *Myripristis* spp. at either stage (Fig. 2). Lastly, the highest densities for each cell type were similar around settlement, irrespective of subfamily, but by adulthood, *S. rubrum* had much lower peak rod densities and higher peak cone and GC densities than *Myripristis* spp.

### Opsin gene expression

Opsin gene expression was analysed across ontogeny in eight species in Holocentridae (four species per subfamily). Phylogenetic reconstruction resolved the class of each opsin gene (Fig. 3) and quantitative opsin gene expression analyses revealed stage-specific expression patterns (Fig. 4, Table 3). At pre-settlement, *S. rubrum* (the only species obtained at this stage) expressed one rod opsin and eight cone opsins: the rod-specific *rh1* (mean ± s.e.m., 78±5% of total opsin gene expression), violet-blue-sensitive *sws2a* (89±9% of single cone opsin gene expression) and *sws2b* (11±9% of single cone opsin gene expression), five green-sensitive *rh2*s (*rh2-1* – *rh2-5*: 7±1 – 34±3% of double cone opsin gene expression), and red-sensitive *lws* (3±2% of double cone opsin gene expression).

**Table 3.** Opsin gene expression in holocentrids over ontogeny. Proportional opsin gene expression in Holocentrinae and Myripristinae (see legend of Fig. 4 for details of species and number of individuals). Values are mean ± s.e.m. given as percentage of total opsin gene expression (*rh1*), single cone opsin gene expression (*sws2a, sws2b*) or double cone opsin gene expression (*rh2, lws*). *rh1*, rhodopsin-like middle-wavelength sensitive 1 (rod opsin); *rh2*, rhodopsin-like middle-wavelength sensitive 2; *sws2*, short-wavelength-sensitive 2; *lws*, long-wavelength-sensitive.

| Gene | Pre-settlement larvae – Holocentrinae |  | Settlement larvae - Holocentrinae |  | Settled juveniles – Holocentrinae |  | Adults – Holocentrinae |  |
| --- | --- | --- | --- | --- | --- | --- | --- | --- |
|  | Mean | s.e.m. | Mean | s.e.m. | Mean | s.e.m. | Mean | s.e.m. |
| *rh1* | 77.77 | 5.18 | 97.03 | 2.6 | 92.5 | 0.99 | 97.39 | 0.86 |
| *sws2a* | 89.13 | 8.63 | 94.21 | 3.55 | 96.9 | 1.42 | 100 | 0 |
| *sws2b* | 10.87 | 8.63 | 5.79 | 3.55 | 3.1 | 1.42 | 0 | 0 |
| *rh2-1* | 33.56 | 3.29 | 62.98 | 6.18 | 42.55 | 4.27 | 57.02 | 1.89 |
| *rh2-2* | 23.99 | 0.08 | 36.3 | 6.2 | 35.17 | 2.83 | 42.98 | 1.89 |
| *rh2-3* | 6.59 | 0.77 | 0.46 | 0.28 | 18.81 | 4.59 | 0 | 0 |
| *rh2-4* | 6.83 | 6.28 | 0 | 0 | 3.29 | 2.78 | 0 | 0 |
| *rh2-5* | 26.18 | 4.32 | 0 | 0 | 0.03 | 0.03 | 0 | 0 |
| *lws* | 2.85 | 2.19 | 0.26 | 0.15 | 0.15 | 0.06 | 0 | 0 |
|  |  |  | Settlement larvae – Myripristinae |  |  |  | Adults – Myripristinae |  |
|  |  |  | Mean | s.e.m. |  |  | Mean | s.e.m. |
| *rh1* |  |  | 89.42 | 1.01 |  |  | 99.57 | 0.07 |
| *sws2a* |  |  | 97.57 | 0.92 |  |  | 100 | 0 |
| *sws2b* |  |  | 2.43 | 0.92 |  |  | 0 | 0 |
| *rh2-1* |  |  | 96.38 | 1.13 |  |  | 100 | 0 |
| *rh2-2* |  |  | 1.44 | 0.36 |  |  | 0 | 0 |
| *rh2-3* |  |  | 1.02 | 0.38 |  |  | 0 | 0 |
| *lws* |  |  | 1.13 | 0.49 |  |  | 0 | 0 |

**Fig. 3.**
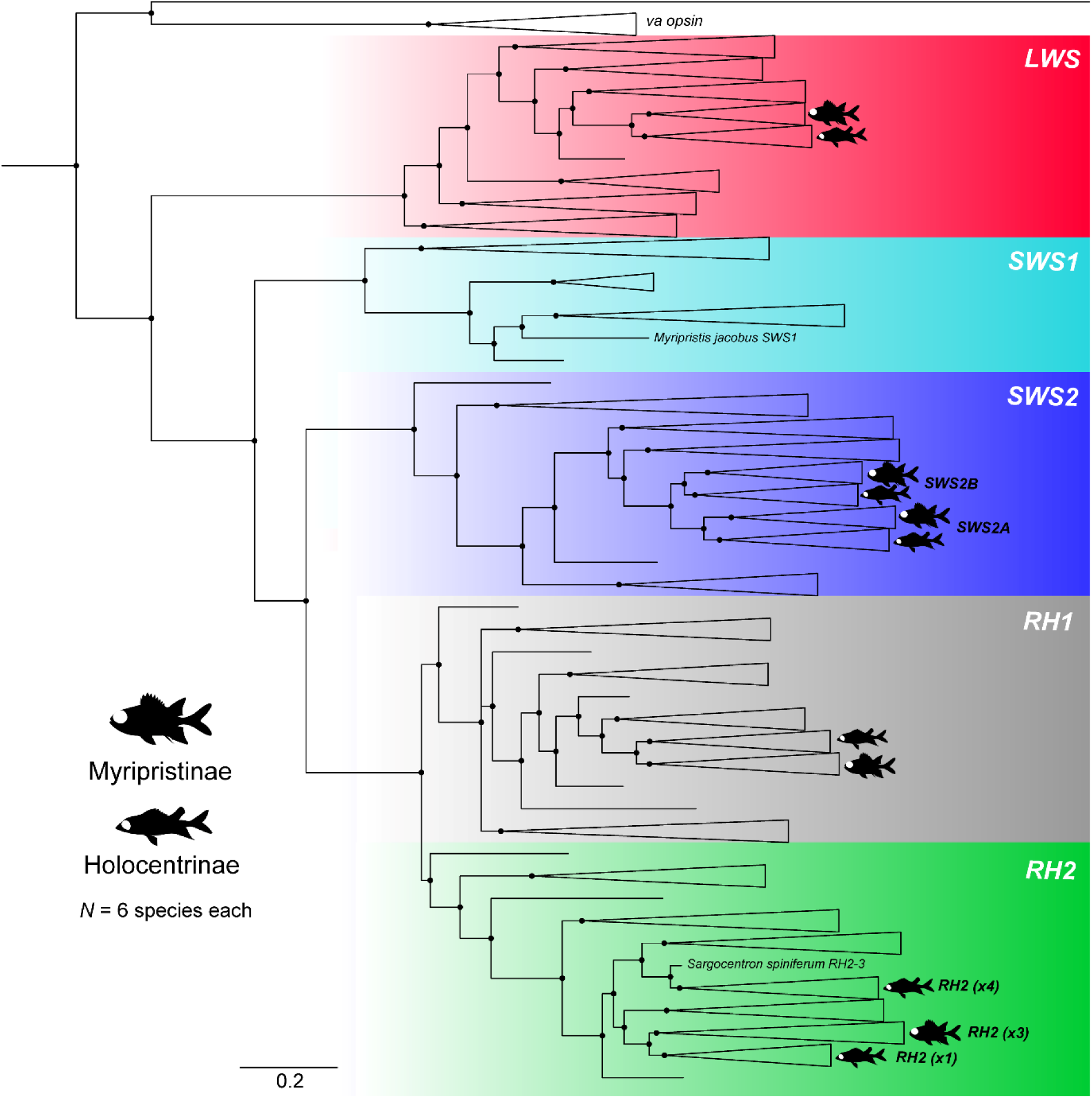
Vertebrate opsin gene phylogeny. Phylogeny including opsin genes from six species per subfamily in Holocentridae denoted as per legend. All expressed genes fell into four out of the five main classes (Bowmaker 2008), while the only *sws1* sequence was derived from a genome (Musilova *et al*. 2019). Numbers in brackets are number of *rh2* paralogs. Black circles denote Bayesian posterior probabilities > 0.8. The scalebar denotes substitutions per site. *rh1*, rhodopsin-like middle-wavelength sensitive 1 (rod opsin); *rh2*, rhodopsin-like middle-wavelength sensitive 2; *sws1*, short-wavelength-sensitive 1; *sws2*, short-wavelength-sensitive 2; *lws*, long-wavelength-sensitive; *va opsin*, vertebrate ancient opsin (outgroup). Accession numbers are provided in Table S1.

**Fig. 4.**
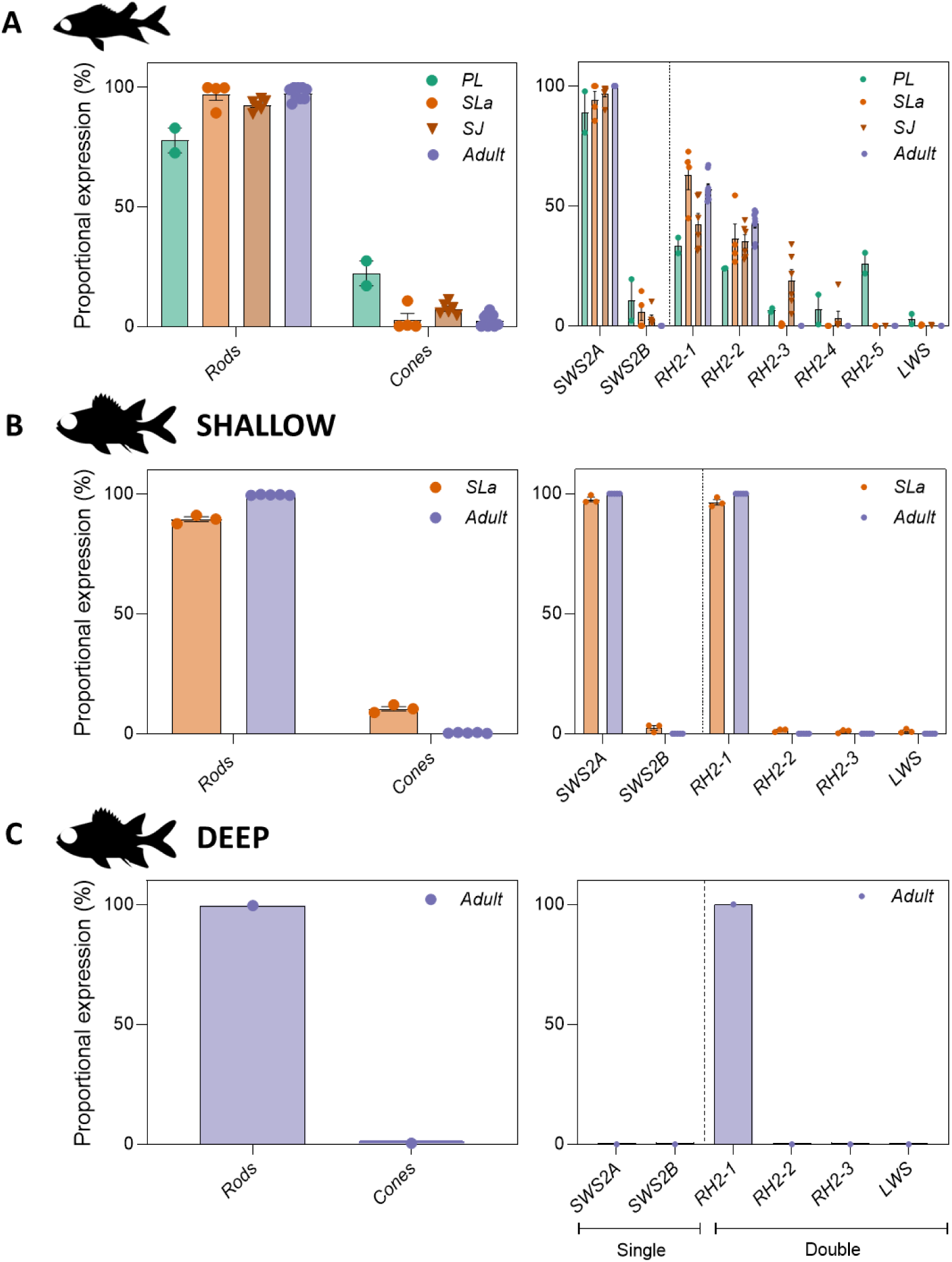
Opsin gene expression in holocentrids over ontogeny. Proportional rod opsin gene expression (given as % of total opsin gene expression) and proportional cone opsin gene expression [given as % of single (*sws’*s) or double (*rh2*s and *lws*) cone opsin gene expression] in (A) Holocentrinae [pre-settlement larvae (*n*=2), settlement larvae (*n*=4), settled juveniles (*n*=6) and adults (*n*=9) from *Sargocentron rubrum, S. punctatissimum, S. cornutum, Neoniphon sammara*], (B) shallow-dwelling Myripristinae [settlement larvae (*n*=3) and adults (*n*=5) from *Myripristis berndti, M. kuntee*, and *M. pralinia*], and (C) deep-dwelling Myripristinae [adults (*n*=1) from *Ostichthys* sp.]. Data are mean ± s.e.m. Green, pre-settlement larvae (PL); light orange, settlement-stage larvae (SLa); dark orange, settled juveniles (SJ); purple, adults. *rh1*, rhodopsin-like middle-wavelength sensitive 1 (rod opsin); *rh2*, rhodopsin-like middle-wavelength sensitive 2; *sws2*, short-wavelength-sensitive 2; *lws*, long-wavelength-sensitive.

At settlement, all species in Holocentridae expressed one rod opsin and six to eight cone opsins: an *rh1* (Holocentrinae: 97±3%, Myripristinae: 89±1%), *sws2a* (Holocentrinae: 94.2±3.6%; Myripristinae: 97.6±1%), *sws2b* (Holocentrinae: 5.8±3.6%; Myripristinae: 2.4±1%), *lws* (Holocentrinae: 0.3±0.16%; Myripristinae: 1.1±0.5%) and several *rh2* paralogs (Holocentrinae: 63±6%; Myripristinae: 96.4±1%; % for the most highly expressed paralog, *rh2-1*) (Fig. 4, Table 3). Most species in Holocentrinae expressed four or five *rh2* paralogs at settlement, and one of these was expressed at low levels (≤0.5%) in every species. All species in Myripristinae expressed three *rh2* paralogs at settlement, with two of these expressed at low levels (≤1.5% per paralog). Between settlement and adulthood, all holocentrids increased rod opsin gene expression, decreased cone opsin gene expression, and stopped expressing three to four cone opsin genes (*sws2b, lws* and 1-2 *rh2*s).

As adults, all shallow-water holocentrids retained some similarities in their opsin gene repertoires, expressing one rod opsin and two to three cone opsins. All species expressed an *rh1* (Holocentrinae: 97.4±0.9%; Myripristinae: 99.6±0.1%) and *sws2a* (100% in both subfamilies) opsin gene, while the variation in *rh2* paralogs remained, with two *rh2* paralogs expressed in Holocentrinae (*rh2-1*: 57±2%, *rh2-2*: 43±2%) and only one in Myripristinae (100%) (Fig. 4, Table 3). Finally, an adult of the deeper-dwelling species from Myripristinae (*Ostichthys* sp.) expressed a simpler opsin gene repertoire than the shallow-water species, with one *rh1* gene (99.6%) and one *rh2* paralog (100%). Overall, opsin gene expression differed between the subfamilies across development, with differences in per-gene expression levels and the number of opsin classes and *rh2* paralogs expressed. Most notably, Myripristinae showed a greater increase in rod opsin gene expression post-settlement than Holocentrinae and expressed one less *rh2* paralog upon maturity.

### Whole-retina differential gene expression

Differential gene expression across the entire retinal transcriptome was examined over development for *S. punctatissimum*. Transcriptomes separated distinctly by life stage in a PCA (Fig. 5A). Whole-transcriptome analyses revealed that a total of 8,395 out of 54,094 transcripts were differentially expressed over ontogeny (adjusted p-value < 0.05). Upon annotation, this total was refined to 4,327 differentially expressed genes (DEGs) (*i*.*e*., 8% of transcripts) which matched to 4,777 gene ontology (GO) terms. Of the DEGs, 1,163 were upregulated in settlement larvae (27%) and 3,164 were upregulated in adults (73%) (Fig. 5B). The top ten differentially expressed genes (DEGs) upregulated in larval retinas were largely involved in cell differentiation/proliferation and cellular structure. Conversely, the top 10 DEGs upregulated in adults were primarily involved in visual perception and aerobic respiration (Table 4).

**Table 4.** Top upregulated genes in *Sargocentron punctatissimum* over ontogeny. Tabular summary of annotations for the top 10 upregulated genes (*i*.*e*., greatest fold change) at settlement larval and adult stages, along with (non-exhaustive) gene function descriptions from QuickGo (Binns *et al*. 2009).

| Settlement larvae |  |  |
| --- | --- | --- |
| Function | Gene ID | Gene name |
| Cell differentiation/proliferation | *prox1* | Prospero homeobox protein 1 |
|  | *dll4* | Delta-like protein 4 |
|  | *fgfp1* | Fibroblast growth factor-binding protein 1 |
|  | *atf5* | Cyclic AMP-dependent transcription factor ATF-5 |
| Basement membrane structure | *co4a1* | Collagen alpha-1(IV) chain |
|  | *co4a2* | Collagen alpha-2(IV) chain (Fragment) |
| Lysosome trafficking | *vps41* | Vacuolar protein sorting-associated protein 41 homolog |
| Cone-specific phototransduction | *arrc* | Arrestin-C |
| Transposition | *lorf2* | LINE-1 retrotransposable element ORF2 protein |
| Angiogenesis | *trbm* | Thrombomodulin |
| Adults |  |  |
| Function | Gene ID | Gene name |
| Visual perception | *opsd* | Rhodopsin |
|  | *opsg* | Green-sensitive opsin |
|  | *opsg2* | Green-sensitive opsin-2 |
|  | *cnrg* | Retinal rod rhodopsin-sensitive cGMP 3',5'-cyclic phosphodiesterase subunit gamma |
| Aerobic respiration | *cox2* | Cytochrome c oxidase subunit 2 |
|  | *cyb* | Cytochrome b |
|  | *cox1* | Cytochrome c oxidase subunit 1 |
|  | *cox3* | Cytochrome c oxidase subunit 3 |
|  | *atp6* | ATP synthase subunit a |
|  | *flvc2* | Feline leukemia virus subgroup C receptor-related protein 2 |

**Fig. 5.**
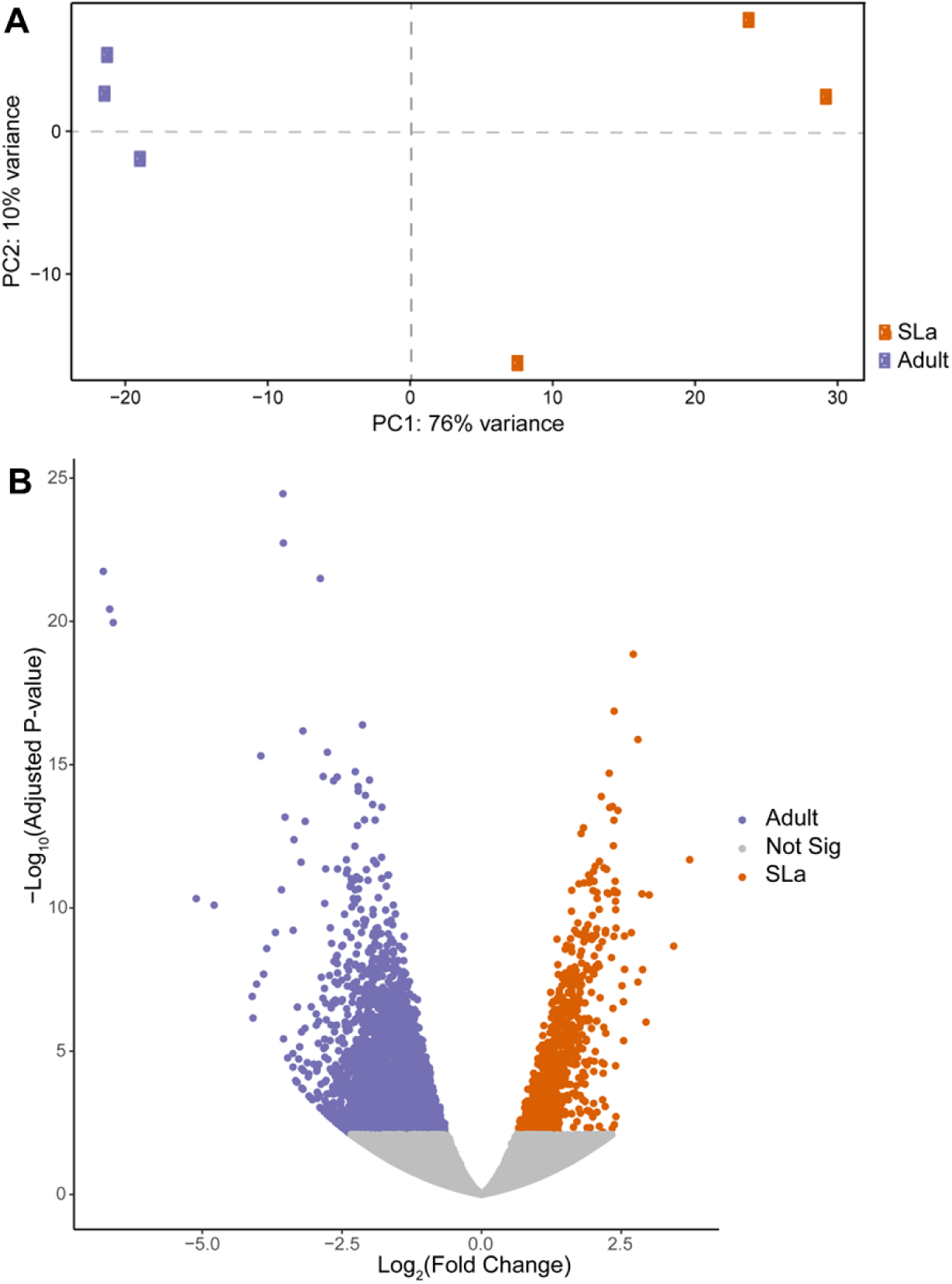
Differential gene expression in *Sargocentron punctatissimum* over ontogeny. (A-B) Differential retinal gene expression between settlement larvae (SLa; *n*=3) and adult (*n*=3) *S. punctatissimum* (Holocentrinae). (A) PCA plot of rlog transformed gene counts showing the variance between individual transcriptomes used for differential gene expression analyses. (B) Volcano plot depicting (log_2_) fold changes in expression of all retinal genes against the (-log10 of the) adjusted p-value. Orange dots with a log fold change > 0 represent transcripts that were significantly higher expressed in settlement larvae, and purple dots with a log fold change < 0 were significantly higher expressed in adults. Grey dots represent transcripts that were not differentially expressed.

Over development, 174 GO terms were found to be enriched (FDR-adjusted p-value < 0.01 and fold enrichment ≥ 2, Table S3). The top three GO terms were related to transcriptional activity: mRNA polyadenylation (GO:0006378), nuclear-transcribed mRNA catabolic process, nonsense-mediated decay (GO:0000184) and RNA polyadenylation (GO:0043631). Many of the other GO terms enriched over development were related to more general cellular processes, such as protein targeting to ER (GO:0045047) and histone modification (GO:0016570). However, analyses also revealed the enrichment of terms relating specifically to visual development: eye morphogenesis (GO:0048592), sensory organ morphogenesis (GO:0090596), eye development (GO:0001654), visual system development (GO:0150063), retinal ganglion cell axon guidance (GO:0031290) and neuron development (GO:0048666).

## Discussion

### Development of the multibank retina

The multibank retina is one of the most common visual specialisations in deep-sea fishes, found in at least 38 families from across the teleost phylogeny (de Busserolles *et al*. 2020; Awaiwanont *et al*. 2001). Based on the few studies on multibank retina development, it appears that rod banks are added as fish grow (Locket 1980; Pankhurst 1987; Frohlich and Wagner 1998; Wagner *et al*. 1998; Omura *et al*. 2003; Taylor *et al*. 2011, 2015), either continually (for mesopelagic fishes and one catadromous elopomorph species, *Anguilla japonica*), or until 20 – 47% of the maximal size is reached (for bathypelagic fishes) (Locket 1980; Pankhurst 1987; Frohlich and Wagner 1998; Omura *et al*. 2003). Similar to bathypelagic fishes, the present study showed that in holocentrids, banks were added as the fish grew, up until they reached 40-60% of their maximal size. However, this may be found to be even earlier if more intermediate sizes were examined. Moreover, most banks were added after holocentrids settled on the reef and transitioned to a dimmer environment. Whether the addition of rod banks was driven by the exposure to dim light is still unknown. However, light environment has consistently been shown to be the dominant driver of visual adaptations in marine fishes (Shand *et al*. 2008; Cortesi *et al*. 2016; Luehrmann *et al*. 2018; Schweikert *et al*. 2018) and thus, represents a convincing possibility.

Further evidence for light environment as a driver of multibank retina development comes from an examination of intraretinal and interspecific variability in bank numbers. This variability has been reported in some adult deep-sea fishes (Locket 1985; Denton and Locket 1989) as well as some adult holocentrids (de Busserolles *et al*. 2021). Similarly, this study showed that the number of banks in adult holocentrids varied with both retinal region and species. However, at earlier stages, rod banking did not vary greatly with either factor. Thus, the holocentrid multibank retina only became specialised later in life once they had adopted a dim light environment. This implies that the multibank adaptation does not become fully active until maturity and/or under dim conditions.

Despite the prevalence of multibank retinas, their function remains a mystery. Two main non-mutually exclusive hypotheses have been proposed: 1) multibank retinas enhance luminous sensitivity (Frohlich and Wagner 1998) and/or, 2) they allow colour vision in dim light (Denton and Locket 1989). Results from this study seem to support both ideas. Support for the sensitivity hypothesis comes from the co-localisation of peak rod:GC convergence and peak rod banking in Myripristinae, suggesting that summation of visual signals is prioritised in their multibank retina. Notably, if the multibank retina solely enhanced sensitivity, a positive correlation between maximum bank number and a species’ depth might be expected, but this is not the case (Fig. 6). Conversely, support for the colour vision hypothesis comes from the co-localisation of peak INL cell densities with peak cone densities at settlement but peak rod densities in adulthood. Given that the INL is the primary layer for opponent processing (Baden and Osorio 2019), this potentially suggests a developmental switch in opponent processing of cone- to rod-derived signals. Although these are intriguing insights, future electrophysiological and behavioural studies are required to confirm the function of the multibank retina throughout ontogeny.

**Fig. 6.**
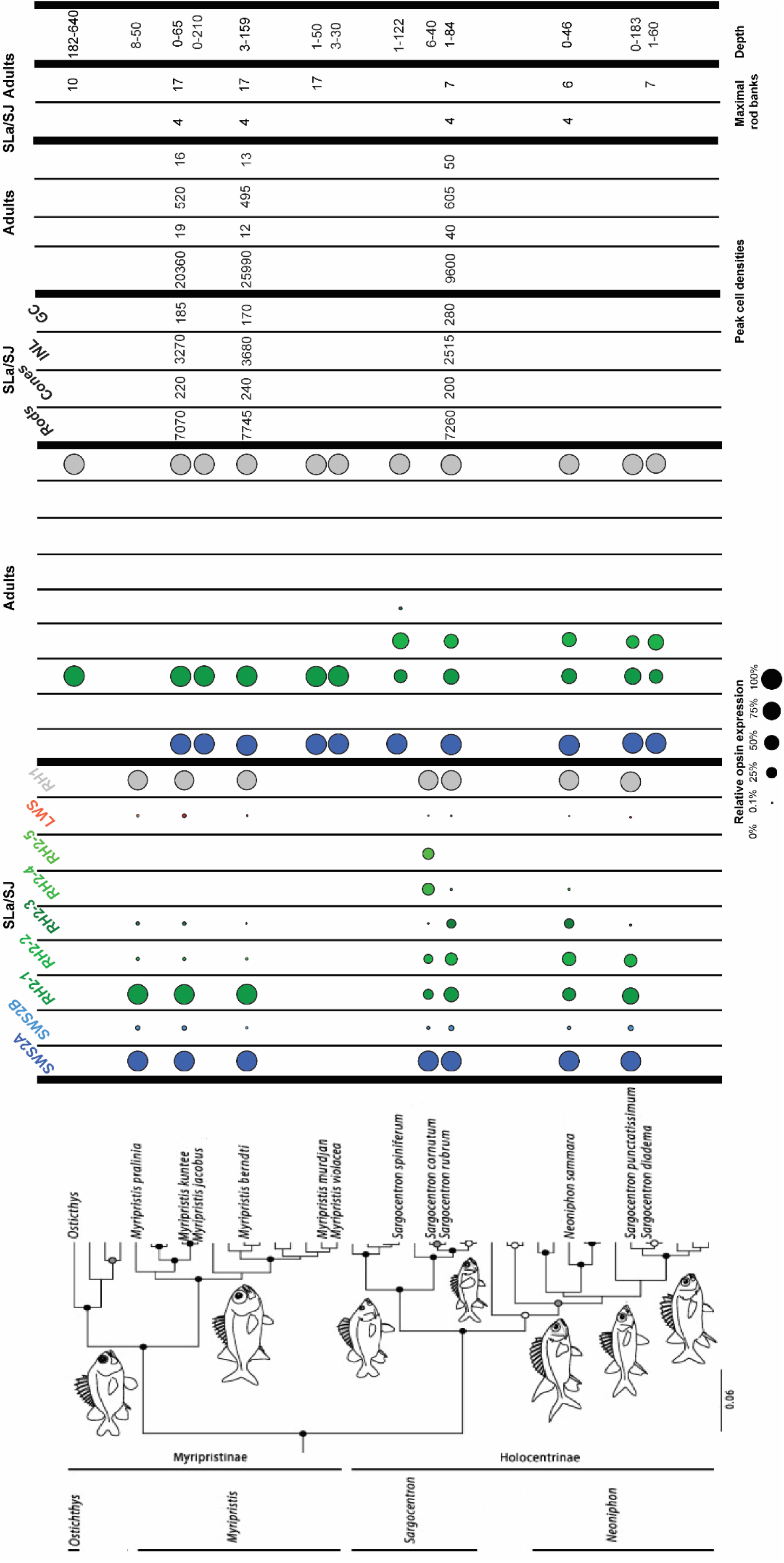
Summary of visual adaptations in holocentrids over development. Opsin gene expression, peak retinal cell densities and maximal rod banking in settlement larvae (SLa) or settled juveniles (SJ) and adults (A) alongside depth at maturity (in metres) are overlaid onto the holocentrid phylogeny. Dots represent the expression of opsin genes in the transcriptome with the size of the dot illustrating relative opsin gene expression as a percentage of total (*rh1*), single cone (*sws2a* and *sws2b*) or double cone (*rh2* and *lws*) opsin gene expression. Peak cell densities (given as cells per 0.01 mm^2^ of retina) and maximal rod banking are the highest densities of each respective cell type and the highest number of banks, respectively, found at the given stage after examining the dorsal, ventral, central, nasal, and temporal retina. Phylogeny adapted from Dornburg *et al*. (2012). Note that *Ostichthys* specimen could not be identified to a species level and so the depth for *Ostichthys kaianus* is shown. The maximal rod banking given for the *Ostichthys* species is an estimation since only ventral sections were available. References for depth: *O. kaianus* (Greenfield *et al*. 2017), *M. pralinia* (Allen 1988), *M. jacobus* (Moore 2015), *M. murdjan* (Lieske 1994), *M. violacea* (Allen 2012), *M. kuntee* (Bacchet *et al*. 2016), *M. berndti* (Allen 2012), *S. spiniferum* (Lieske 1994), *S. cornutum* (Allen 2012), *S. punctatissimum* (Lieske 1994), *S. diadema* (Randall 1998), *S. rubrum* (Randall 1998) and *N. sammara* (Lieske 1994). *rh1*, rhodopsin-like middle-wavelength sensitive 1 (rod opsin); *rh2*, rhodopsin-like middle-wavelength sensitive 2; *sws2*, short-wavelength-sensitive 2; *lws*, long-wavelength-sensitive.

### Retinal cell densities over development

Most teleosts commence life with a pure cone retina, with rods added later (Evans and Fernald 1990; Raymond *et al*. 1995). While most diurnal shallow-water fishes follow this developmental trajectory (Blaxter and Staines 1970), it may be adjusted when faced with different ecological demands. For example, deep-sea or nocturnal fishes show more rapid and pronounced increases in rod densities and decreases in cone densities over development (Shand 1997; Locket 1980; Pankhurst 1987; Bozzano *et al*. 2007). In line with their ecology, holocentrids followed a nocturnal pattern, rapidly decreasing cone densities and increasing rod densities. These retinal changes were particularly pronounced post-settlement correlating with the timing at which holocentrids are thought to become nocturnal (Shand 1994b). Moreover, the extent of developmental changes differed between the two subfamilies. At settlement, both subfamilies had similar visual systems. However, in adults, higher rod densities and lower cone densities were found in Myripristinae compared to Holocentrinae, similar to findings from retinal wholemounts (de Busserolles *et al*. 2021). Holocentrinae retained more of their photopic visual system, the reason for which requires further studies on their day-time activities. In summary, the holocentrid visual system is remodelled at the cellular level over development to suit their nocturnal lifestyle, while still maintaining some adaptation for daytime activity.

Shallow-water holocentrids are thought to have emerged from a deep-water existence and some of their extant relatives are still found at greater depths, down to 640 metres (Yokoyama *et al*. 2008; Greenfield *et al*. 2017). Given this phylogenetic connection to the deep-sea, it is not surprising that some aspects of their visual development were comparable to deep-sea fishes while others more closely resembled shallow-water fishes. In terms of the cones, a steep decline in densities was evident during development and the adult population was mainly composed of double cones, similar to what is found in some deep-sea fishes (Boehlert 1979; Munk 1990; de Busserolles *et al*. 2021). However, holocentrids retained cones in all retinal regions throughout life while these are often lost at early developmental stages (Bozzano *et al*. 2007) or become restricted to certain retinal regions (Munk 1990) in some deep-sea fishes. With respect to rods, adult holocentrids (particularly in Myripristinae) possessed peak densities that rival those of some deep-sea fishes (*e*.*g*., *Myripristis berndti*: ∼2.6 million rods mm^-2^ vs. *Myctophum brachygnathum*: ∼2 million rods mm^-2^, and *Hoplostethus atlanticus*: ∼1.7 million rods mm^-2^) (Pankhurst 1987; de Busserolles 2013) and their maximal rod:GC summation even exceeds that of many deep-sea species (*e*.*g*., *Myripristis berndti:* 3800:1 vs. *Lampanyctodes* spp.: 2000:1, and *Chauliodus sloani*: 200:1) (Locket 1980; Pankhurst 1987). Finally, the developmental decrease in GC densities in holocentrids is intermediate compared to the very steep decrease observed in deep-sea fishes (Locket 1980; Pankhurst 1987) and the more subtle change found in diurnal shallow-water species (Johns and Easter 1975; Shand *et al*. 2000).

### Ontogenetic shifts in retinal specialisations

Retinal specialisations in teleosts usually reflect ecological demands (Collin and Pettigrew 1988a, 1988b; Luehrmann *et al*. 2020; de Busserolles *et al*. 2014b; Collin 2008) and, accordingly, have been shown to shift during ontogeny (Shand *et al*. 2000; Tettamanti *et al*. 2019). This is also the case in the holocentrids. At settlement, all species had similar retinal specialisations. The region with greater acuity (*i*.*e*., highest GC densities) and better adaptation for bright light vision (*i*.*e*., highest cone densities) was found in the temporal retina. This area surveys the visual field immediately in front of the fish, which may help the larvae to see their small zooplankton prey in the brightly lit surface layers of the ocean (Kawamura *et al*. 1984; Shand *et al*. 2000). On the other hand, larval holocentrids showed the highest sensitivity (*i*.*e*., highest rod densities and rod:GC ratio) in the dorsal retina, which surveys the visual field beneath the fish and may be used to spot predators coming from the dimmer waters below (Collin and Partridge 1996).

After holocentrids have settled on the reef and adopted their nocturnal lifestyle, their retinal specialisations shift accordingly. In adults, the regions with the highest acuity (*i*.*e*., highest GC densities) were located temporally in Holocentrinae (*i*.*e*., looking forward) and ventro-temporally in Myripristinae (*i*.*e*., looking forward and upwards). These specialisations are likely linked to their nocturnal feeding ecologies. As benthic feeders, prey is viewed in front of Holocentrinae, while Myripristinae feed in the water column where food items occur in front of and above fishes [also see: de Busserolles *et al*. (2021)]. The regions which are best adapted for sensitivity (*i*.*e*., highest rod densities) overlapped with the regions of higher acuity in both subfamilies (Holocentrinae: central and temporal; Myripristinae: ventral) and so may also facilitate nocturnal feeding. Finally, the regions with the greatest adaptation for bright light vision (*i*.*e*., highest cone densities) were located temporally in Holocentrinae and centrally in Myripristinae, surveying the area in front of or lateral to the fish, respectively. Little is known about the day-time activities of holocentrids; however, these areas may be linked to social interactions and identification of safe havens for refuge during the day (Winn *et al*. 1964; Carlson and Bass 2000).

### Opsin gene expression over development

Teleosts are known to tune their spectral sensitivities to the light environment over development by changing their relative opsin gene expression levels and/or by switching between gene classes or copies (Shand *et al*. 2002; Cheng and Flamarique 2007; Savelli *et al*. 2018; Luehrmann *et al*. 2018; Musilova *et al*. 2019). Holocentrids are no exception (Musilova *et al*. 2019; de Busserolles *et al*. 2021; Musilova and Cortesi 2021). Developmental changes in holocentrid opsin gene expression correlated well with their switch to a nocturnal, reef-dwelling lifestyle. For example, holocentrids increased their relative *rh1* expression by nearly 20% over development. Furthermore, they stopped expressing most of their cone opsin genes, only retaining cone opsins sensitive to the mid-range (blue – green; *sws2a* and *rh2*) wavelengths that dominate the reef at night. This is in contrast with changes in most diurnal fishes, which do not reduce the number of cone opsins expressed over ontogeny (Takechi and Kawamura 2005; Cheng and Flamarique 2007; Shand *et al*. 2008; Cortesi *et al*. 2016; Tettamanti *et al*. 2019; Chang *et al*. 2020).

Among the cone opsin genes, ontogenetic changes in *rh2* expression are particularly interesting in holocentrids. With eight copies, holocentrids have the highest number of genomic *rh2* paralogs of any teleost species (Musilova *et al*. 2019). Our results show that most of these *rh2* paralogs, along with several other cone opsin genes, were only expressed at early life stages. Moreover, some of these genes (*e*.*g*., *lws*) were expressed at low levels that may not be functionally relevant to vision but may instead represent the remnants of an opsin cascade that regulates photoreceptor development, as observed for salmonoid and cyprinid fishes (Raymond *et al*. 1995; Stenkamp *et al*. 1996; Takechi and Kawamura 2005; Cheng *et al*. 2007). However, *rh2* was the only cone opsin gene subclass expressed in all stages/species in Holocentridae (Fig. 6) and this subclass is sensitive to wavelengths present in the light environment of these fishes throughout life. Thus, the different *rh2* paralogs may serve a visual purpose, allowing the fish to switch between opsin gene palettes during ontogeny for more precise control over spectral tuning (known as subfunctionalisation), similar to findings in cichlids (Spady *et al*. 2006).

Although differences in opsin gene expression in teleosts are often explained by the light environment and species-specific ecologies, phylogenetic forces also exert control (Tettamanti *et al*. 2019; Carleton *et al*. 2008; Stieb *et al*. 2016). This may also be the case in holocentrids. Although shallow-dwelling species share a similar light environment at every life stage, the two subfamilies only showed similar opsin gene expression early in life. As adults, shallow-dwelling Myripristinae expressed fewer cone opsin genes and higher *rh1* levels than Holocentrinae. This more extreme adaptation for dim-light vision in shallow-dwelling *Myripristis* spp. may be because they are more closely related to deep-dwelling *Ostichthys* spp., resulting in greater similarity to their deep-water relatives. Notably, this potential phylogenetic inertia did not seem to completely negate the influence of ecological drivers. As such, fewer cone opsins were expressed in an adult from the deeper-dwelling genus *Ostichthys* (Myripristinae) compared to shallow-dwelling Myripristinae representatives, aligning well with the depth-related narrowing of spectral sensitivities observed in other teleosts (Schweikert *et al*. 2018).

### Retinal gene expression over development

Around 8% of annotated retinal transcripts were differentially expressed over development in holocentrids. Although whole-retina changes in gene expression have not been studied in teleosts that undergo major ecological shifts, some insights can be gained from other taxa. For example, butterflies show developmental changes in visual gene expression alongside changes in eye structure and therefore, some of the stage-specific expression differences were attributed to cellular composition (Ernst and Westerman 2021). Similarly, since the holocentrid retina underwent cellular remodelling over development, it is likely that some of the expression changes simply facilitate these structural changes. Indeed, several retinal genes which were highly upregulated in settlement larvae are involved in cell differentiation and proliferation (*e*.*g*., *rx3* and *otx2*) (Loosli *et al*. 2003; Yamamoto *et al*. 2020).

Interestingly, similar to findings in frogs (Valero *et al*. 2017; Schott *et al*. 2021), several of the differentially expressed genes were correlated with ecological changes over development. For example, genes involved in cone- or rod-based photoreception were upregulated in larval or adult holocentrids, respectively, aligning with their switch from bright to dim environments. Also, the most highly upregulated genes in adults were mainly involved in photoreception. This is congruent with the higher photoreceptor densities of adults and implies that adults have a higher investment in vision than larvae. Overall, in holocentrids, developmental changes to retinal gene expression align well with the retinal structure and ecological demands of each life stage.

## Conclusion

The holocentrid visual system adapted to life in dim light over ontogeny. At the morphological level, they increased rod banks from 1-2 to 5-17, adopted a rod-dominated retina and increased visual summation. At the molecular level, they increased rod opsin gene expression, narrowed the cone opsin gene expression repertoire from 8 to 1-4 cone opsins, and shifted from enrichment of cell differentiation/proliferation to phototransduction. Despite the early emergence of the multibank retina, substantial topographic specialisations in bank number were only evident after the transition to a dimmer environment. Together, this suggests that ecology drives visual development in Holocentridae. However, subfamily-specific differences in the degree of scotopic specialisation emerged over development (*i*.*e*., more rod banks, higher rod densities, greater summation, and higher rod opsin gene expression in Myripristinae) and these were correlated with phylogenetic relatedness to deep-water representatives rather than ecology. This suggests that the development of the holocentrid retina may also be somewhat driven by phylogeny. Future studies on visual development in other nocturnal reef fishes as well as other marine fish families with both shallow- and deep-water forms, such as Anomalopidae (flashlight fishes) and Engraulidae (anchovies), may provide further insights into the ecological and phylogenetic drivers of the development of dim-light vision.

## Acknowledgements

We acknowledge the Dingaal, Ngurrumungu and Thanhil peoples as traditional owners of the lands and waters of the Lizard Island region from where specimens were collected. We also acknowledge the traditional owners of the land on which the University of Queensland is situated, the Turrbal/Jagera people, where this work was done. We would like to thank the staff at Lizard Island Research Station (LIRS), Dr Anne Hoggett and Dr Lyle Vail, as well as the staff at the Centre of Island Research and Environmental Observatory (CRIOBE) for support during field work. We also thank Cairns Marine for supplying animals. We thank Robert Sullivan from the Queensland Brain Institute (QBI) Histology Facility, Richard Webb and Robyn Chapman Webb from the Australian Microscopy & Microanalysis Research Facility at the Centre for Microscopy and Microanalysis (CMM), and Rumelo Amor from the QBI Advanced Microscopy facility for scientific and technical support and advice. We thank Janette Edson from the QBI Genomics Facility and staff at Novogene Co., Ltd for library preparation and transcriptome sequencing. Finally, we would like to thank Professor Mark McCormick and his team for generously lending us their light traps and assisting with some animal collections.

## Data availability

Data will be made available via the University of Queensland’s eSpace at publication.

